# RNA-Bloom provides lightweight reference-free transcriptome assembly for single cells

**DOI:** 10.1101/701607

**Authors:** Ka Ming Nip, Readman Chiu, Chen Yang, Justin Chu, Hamid Mohamadi, René L. Warren, Inanc Birol

## Abstract

We present RNA-Bloom, a *de novo* RNA-seq assembly algorithm that leverages the rich information content in single-cell transcriptome sequencing (scRNA-seq) data to reconstruct cell-specific isoforms. We benchmark RNA-Bloom’s performance against leading bulk RNA-seq assembly approaches, and illustrate its utility in detecting cell-specific gene fusion events using sequencing data from HiSeq-4000 and BGISEQ-500 platforms. We expect RNA-Bloom to boost the utility of scRNA-seq data, expanding what is informatically accessible now.

---

The early stages of single-cell RNA sequencing (scRNA-seq) data analyses have been primarily limited to gene expression quantification and individual splice-junction detection^1^. Isoform-level analyses for single cells remain scarce^2^. Although long-read sequencing technologies excel in capturing near full-length transcript sequences, which are ideal for isoform-level analyses^2^, they have a significantly higher error-rate, limited sequencing depth, and higher input requirement compared to short-read sequencing technologies^3^. Short-read sequencing prepared with transcript-end capture protocols^4-6^ are scalable to nearly one million cells, but they exhibit strong transcript-end bias, prohibiting the reconstruction of alternative isoforms. Lifting this bias, there are other short read sequencing protocols^7, 8^ that offer full-length transcript sequencing, thus permitting both expression quantification and isoform structure analysis. However, to leverage the rich information in the data generated by the latter protocols new bioinformatics approaches are required. The current scRNA-seq assembly methods are primarily intended for specific gene targets^9-11^, and *de novo* assembly methods for bulk RNA-seq do not effectively accommodate uneven transcript coverage and amplified background noise in scRNA-seq data. In principle, pooling reads from multiple cells can introduce reads to low-coverage and noisy regions of transcripts, thus closing coverage gaps and increasing the signal-to-noise ratio (**Supplementary Fig. 1**). Naively pooling reads from multiple cells would obscure the cell-specificity of the assembly process and would require much more memory than assembling sequencing data for each individual cell. Therefore, a resource-efficient scRNA-seq *de novo* assembly method that maintains high cell-specificity of reconstructed transcripts is needed.

Leading *de novo* transcriptome assembly methods, such as Trans-ABySS^12^, follow the de Bruijn graph (DBG) assembly paradigm. While these methods use a hash table data structure to store the DBG in memory, recent genome assembly methods^13, 14^ showed how memory requirements may be reduced by adopting succinct data structures, such as Bloom filters^15^, for compact *k*-mer storage to representing an implicit DBG. Adapting this strategy for RNA-seq assembly is an attractive proposition for reducing memory consumption. Further, short-read sequencing platforms are often used to produce paired-end reads, which can be valuable in reconstructing longer transcript sequences, and existing transcriptome assembly methods rely on alignment of these paired-end reads against assembled sequences to do that. Read alignments are computationally costly, especially when performed against a non-static target, such as earlier stages of an assembly. We note that, recent RNA-seq quantification tools, such as kalisto^16^, reduce runtime by replacing alignment with pseudoalignment. Borrowing the strategy of substituting read alignment with a lightweight alternative should also benefit runtimes when assembling transcriptomes.

These strategies are implemented in our scRNA-seq assembly method, RNA-Bloom. The method leverages sequencing content from multiple cells for improved transcript reconstruction of individual cells. It follows the DBG assembly paradigm but uses Bloom filters to represent graphs for efficient in-memory storage of *k*-mers as well as *k*-mer counts (Online Methods). It utilizes paired distant *k*-mers derived from reads and reconstructed fragments as a fast alternative to read alignments (Online Methods).

RNA-Bloom consists of three stages: (1) shared DBG construction, (2) fragment sequence reconstruction, and (3) transcript sequence reconstruction (**Fig. 1** and Online Methods, Algorithm 1). In stage 1, an implicit DBG is constructed using the *k*-mers from all input reads of all cells. In stage 2, fragment sequences of each cell are reconstructed by connecting the cell’s read pairs using the DBG from stage 1. To maintain the cell-specificity of the transcripts to be reconstructed in stage 3, a new DBG for each cell is created solely using *k*-mers from the cell’s reconstructed fragments following stage 2. In stage 3, using the cell-specific DBGs, fragments are extended outward in both directions to reconstruct transcript sequences.

**Figure 1.**
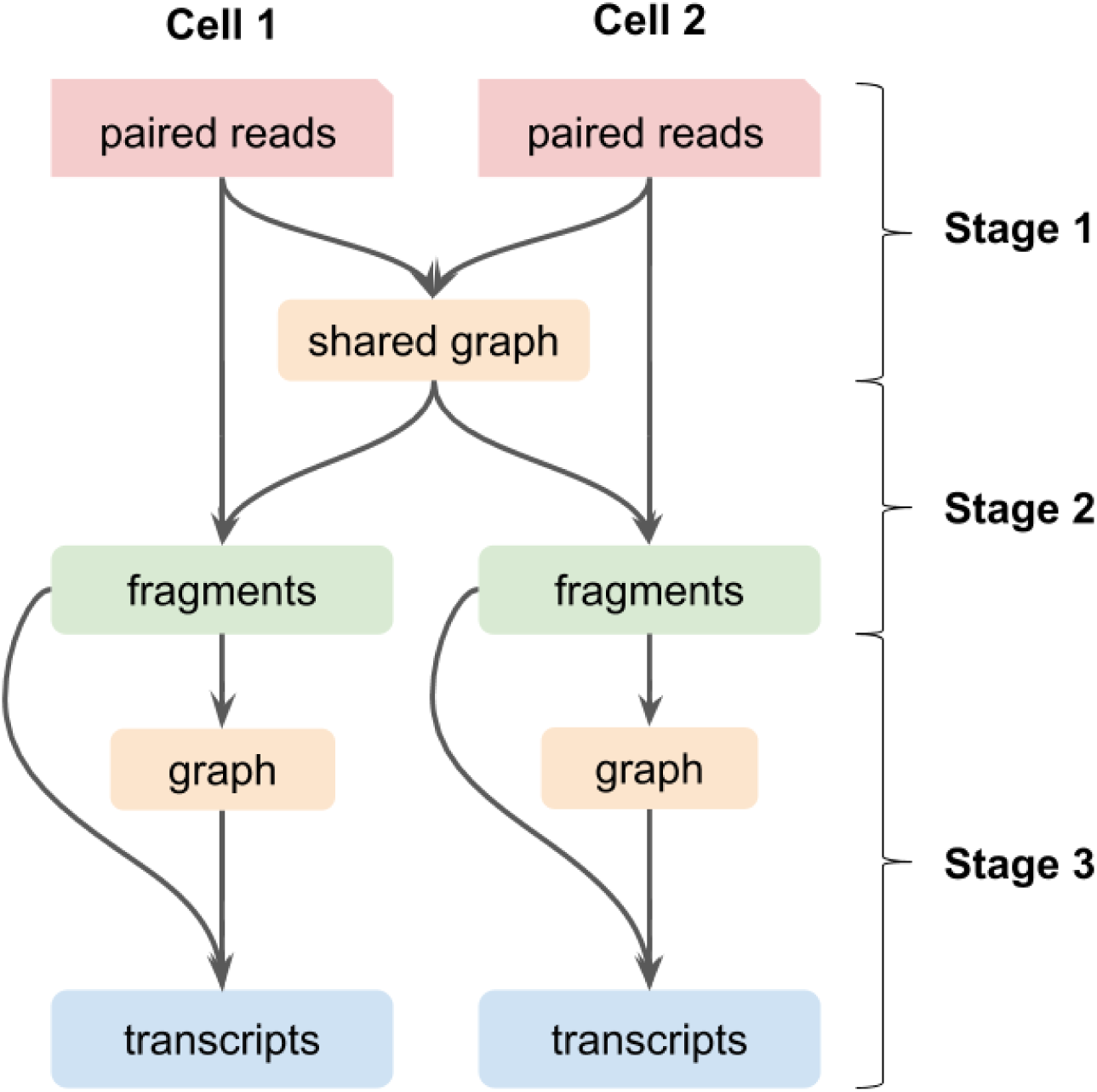
Pooled assembly of scRNA-seq data in RNA-Bloom, illustrated for two cells. RNA-Bloom consists of three stages: (1) construction of a shared de Bruijn graph using reads from all cells, (2) reconstruction of fragments from read pairs of individual cells, and (3) reconstruction of transcripts of individual cells.

To benchmark RNA-Bloom, we compared it against three state-of-the-art *de novo* assemblers for bulk RNA-seq: Trans-ABySS^12^, Trinity^17^, and rnaSPAdes^18^. We examined the computing performance and assembly quality of the four assembly tools on a simulated scRNA-seq dataset of 168 mouse embryonic stem cells (total of 672 million paired-end 100-bp reads). In our tests, RNA-Bloom had the best mean reconstruction of 3,181 (*SD*=498) full-length isoforms per cell, and it assembled on average 30.00% more full-length isoforms than the next best method, Trinity (**Supplementary Fig. 2**). RNA-Bloom maintained a mean proportion of misassemblies (1.69%, *SD*=0.38%), and is comparable to rnaSPAdes (1.43%, *SD*=0.37%) and Trans-ABySS (1.93%, *SD*=0.35%) in its performance, while Trinity had the highest mean proportion of misassemblies (4.08%, *SD*=0.82%) (**Fig. 2c**).

**Figure 2.**
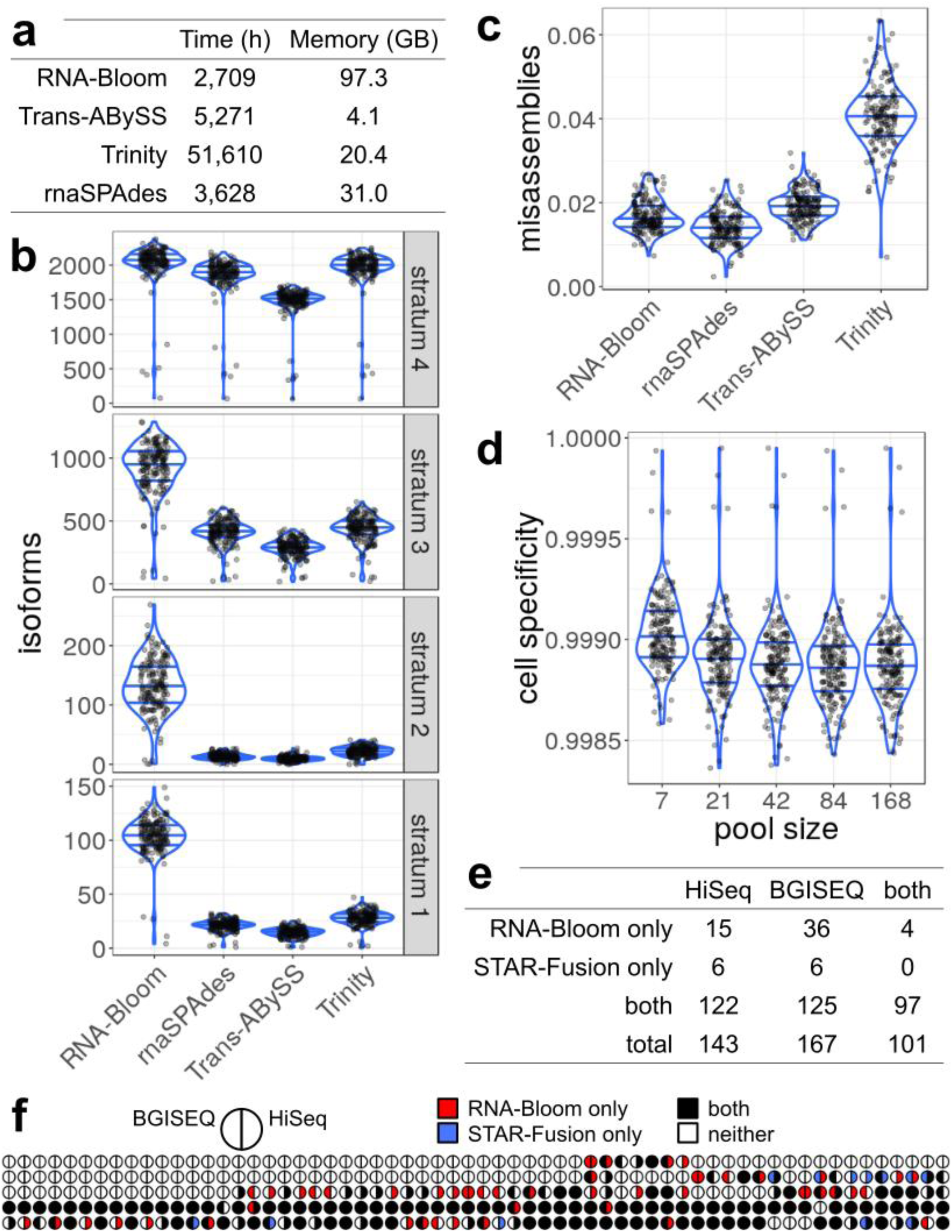
Computing performance and assembly quality of RNA-Bloom and other methods. **(a)** Total run time and peak memory usage of RNA-Bloom, rnaSPAdes, Trans-ABySS, and Trinity. Assemblies were performed on a simulated dataset of 168 mouse embryonic stem cells (4 million paired-end 100-bp reads per cell, total of 672 million reads). All tools were configured to use 48 threads. **(b)** Number of full-length isoforms stratified by the expression level quartiles in the simulated data. Stratums from top to bottom correspond to highest expression to lowest expression. **(c)** Number of misassemblies normalized by the number of full-length isoforms reconstructed. **(d)** Assembly cell-specificity for increasing number of cells in the pool. The pool sizes are selected to ensure that each sub-pool is derived from single pools at larger sizes. **(e)** Total counts for five selected gene fusions detected in 62 cells sequenced with Illumina HiSeq 4000 and BGISEQ-500. **(f)** Dot plot showing the distribution of five selected fusions (rows) across 62 cells (columns). The fusions arranged from top to bottom are: *ABL1:BCR, C16orf87:ORC6, NUP214:XKR3, BAG6:SLC44A4*, and *CBX5:GTSF1*. Colors of half-circles indicate the detection by software tools and sequencing platforms for each cell as indicated on the figure.

We next examined the reconstruction of simulated isoforms at different expression levels. We split the set of all simulated isoforms into four strata based on the quartiles of the expression levels from all cells (Online Methods). We observed RNA-Bloom to have the highest reconstruction figures for all expression strata in all cells, while Trinity was a close second only when the expression levels are high (**Fig. 2b, stratum 4**). All assemblers reconstructed the most full-length isoforms in the highest expression stratum, likely because highly expressed transcripts are represented by more reads, making them easier to assemble. However, the proportional difference in mean reconstruction between RNA-Bloom and bulk RNA-seq assemblers substantially increased in the lower expression strata (1.04x in stratum 4 vs 3.73x in stratum 1). This illustrates that RNA-Bloom’s pooled assembly strategy is working effectively despite potential coverage gaps and amplified noise, which tend to have larger detrimental effect on the reconstruction of low-expressed transcripts due to insufficient good quality reads.

We then examined the effect of the number of cells on RNA-Bloom’s assembly cell-specificity (**Supplementary Note 2**) by down-sampling to smaller pool sizes. We found that the average cell-specificity of RNA-Bloom decreased slightly from 99.90% (*SD*=0.02%) to 99.89% (*SD*=0.02%) as the pool size increased from seven cells to 168 cells (**Fig. 2d**).

RNA-Bloom outperformed all other methods in total runtime. It used only 74.7% of total runtime of the best bulk RNA-seq assembly methods (2,709 CPU hours vs 3,628 CPU hours for rnaSPAdes) (**Fig. 2a**). Trinity had the longest total runtime (51,611 CPU hours), which is 19 times that of RNA-Bloom.

We also explored the scalability of RNA-Bloom using a dataset of 260 mouse embryonic stem cells with a total of 3.94 billion paired-end 100-bp reads^19^. In this dataset, each cell was deeply sequenced to over 9 million reads on average. Using 48 CPUs and 120 GB of memory, RNA-Bloom assembled the dataset in 11 days 21 hours 54 minutes.

To demonstrate RNA-Bloom’s ability to reconstruct unannotated transcripts, we used an scRNA-seq dataset describing the myelogenous leukemia cell line K562 cells (Online Methods), which are known to have five gene fusions (**Supplementary Table 1**). This dataset comprised sequencing data generated with two high-throughput sequencing platforms, Illumina HiSeq 4000 and BGISEQ-500. We used PAVFinder^20^ to detect fusions from single cell transcriptomes assembled by RNA-Bloom. We compared our assembly-first detection method against an alignment-first detection method, STAR-Fusion^21^, which relies on mapping reads against the reference genome. Overall, RNA-Bloom reconstructed 15% more fusion transcripts than those detected by STAR-Fusion (**Fig. 2e**). Although the difference is not significant (Chi-squared test, N=310, *p*=0.242), this illustrates that reference-free assembly methods can be effective in transcript discovery, even when a high-quality reference genome is available. Interestingly, both methods together detected 16.8% more fusions in BGISEQ than HiSeq (Chi-squared test, N=185, *p*=0.0337). This difference is probably a result of the larger number of reads in the BGISEQ libraries. RNA-Bloom reconstructed 22.9% more fusion transcripts in BGISEQ libraries than in HiSeq libraries, but the difference is not statistically significant (Chi-squared test, N=170, *p*=0.202).

New sequencing technologies that interrogate single cell transcriptomes provide information on gene regulation at an unprecedented detail. Whereas transcript-end capture protocols are primarily used to measure expression levels, full-length sequencing protocols have richer information content for isoform structures in single cells. However, to realize the full potential of the full-length sequencing protocols, specialized bioinformatics tools are needed. Here we present RNA-Bloom, a *de novo* single-cell RNA-seq assembly algorithm to address this need. As a scalable assembler demonstrated to work for nearly four billion paired-end reads, RNA-Bloom unlocks the possibility of cataloguing cell types at the isoform level in large datasets. The software is implemented in the Java programming language, and distributed under GPLv3 license. It is available for download in source code and as a pre-built executable JAR file at https://github.com/bcgsc/RNA-Bloom.

## METHODS

Methods and associated references and data accession codes are available in the online version of the paper.

## ACKNOWLEDGEMENTS

This work was supported by Genome Canada and Genome BC [243FOR, 281ANV]; the National Institutes of Health [2R01HG007182-04A1]; and the Natural Sciences and Engineering Research Council of Canada (NSERC). The content of this work is solely the responsibility of the authors, and does not necessarily represent the official views of the National Institutes of Health or other funding organizations.

## AUTHOR CONTRIBUTIONS

K.M.N. and I.B. designed the method and algorithms. K.M.N., J.C., H.M., and I.B. designed the Bloom filter data structures. K.M.N. implemented the RNA-Bloom software. K.M.N., R.C., and C.Y. conducted the benchmarking experiments. K.M.N., R.C., C.Y, R.L.W, and I.B. analyzed the results. All authors wrote the manuscript.

## ONLINE METHODS

### Algorithm 1.

RNA-Bloom reference-free assembly algorithm for scRNA-seq. The overall procedure for each stage of RNA-Bloom is illustrated as pseudocode.

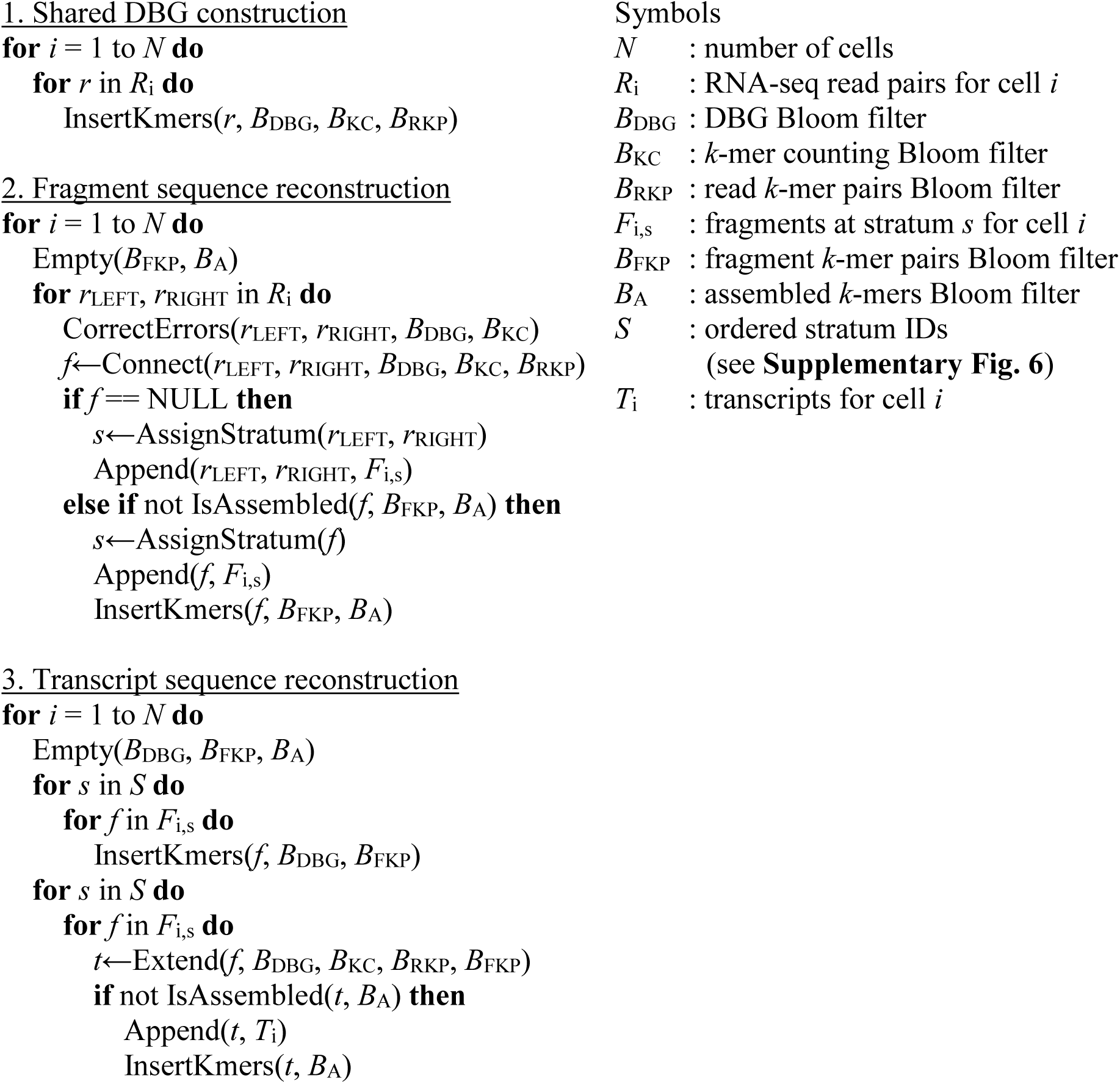

### Bloom filter data structure

The hash functions for Bloom filters in RNA-Bloom were implemented based on the nucleotide hashing algorithm, ntHash^22^. The number of unique *k*-mers and the user-defined false positive rate (FPR) for Bloom filters are used to determine the size for each Bloom filter. RNA-Bloom has the option to run ntCard^23^ to quickly estimate the number of unique *k*-mers, provided ntCard is already installed in the user’s computing environment. If the number of unique *k*-mers is not specified or ntCard is not available, then the total size of Bloom filters is configured in proportion to the total file size of all input reads files. In our experience, a Bloom filter FPR of 0.5∼1.0% provides a relatively good trade-off between memory usage and assembly quality.

### Shared de Bruijn graph construction

The shared DBG is represented by three separate Bloom filters: (1) DBG Bloom filter, (2) *k*-mer counting Bloom filter, and (3) read *k*-mer pairs Bloom filter. The DBG Bloom filter is a bit-array, and it provides an implicit representation of the DBG for the *k*-mers in all cells. The *k*-mer counting Bloom filter provides a compact non-exact storage of *k*-mer counts, and it is implemented based on the 8-bit minifloat byte-array data structure introduced previously^24^. To minimize the effect of false-positives in the Bloom filters, a *k*-mer is deemed present in the dataset only if it is found in the DBG Bloom filter, and it has a non-zero count in the *k*-mer counting Bloom filter. The read *k*-mer pairs Bloom filter stores pairs of distant *k*-mers at a fixed distance along each read (**Supplementary Fig. 3**). These *k*-mer pairs are essentially sparse representations of individual reads, and they are useful in guiding graph traversal in later stages. The shared DBG is used throughout fragment reconstruction of individual cells.

### Fragment sequence reconstruction

After the shared DBG has been constructed, the process of fragment sequence reconstruction is performed separately for individual cells. For each read pair in each cell, mismatch and indel errors are identified and corrected based on *k*-mer counts in a procedure similar to the RNA-seq error correction method, Rcorrector^25^ (**Supplementary Fig. 4**). After error correction, the read pair is connected by extending each constituent read towards its mate to reconstruct the underlying fragment sequence. Each read is extended by searching for neighbors in the shared DBG. When the current extension reaches a branching point in the graph, the unambiguous extension of each branch is assigned a score based on its median *k*-mer count and the number of read *k*-mer pairs spanning across the branching point (**Supplementary Fig. 5** and **Supplementary Note 1**). Since longer extensions tend to have more supporting *k*-mer pairs, the score is normalized by the length of the extension. The branch with the highest score is added to the current extension from the read. This extension routine is initially depth-bounded by a permissive default threshold (default = 1,000 bp) for each cell. After the first N read pairs (default = 1,000) of the cell have been evaluated, the depth threshold is re-adjusted to 1.5-fold of the interquartile range of reconstructed fragment lengths. This threshold limits the depth of graph traversal to ensure fast overall assembly runtime and prevents spurious connections of reads. Extension is first attempted from the left read towards the right and a second attempt is made from the right read towards the left when the first attempt fails. Extension from each direction terminates if the paired reads are connected or the extension has reached either a dead-end or the depth threshold.

Each reconstructed fragment is checked for consistency with input reads by scanning for overlapping read *k*-mer pairs. If the reconstructed fragment is consistent with read *k*-mer pairs, more distant pairs of *k*-mers at a fixed distance within the reconstructed fragment are stored in the fragment *k*-mer pairs Bloom filter (**Supplementary Fig. 3**). The distance between the paired *k*-mers are set to the first quartile of reconstructed fragment lengths from the first N read pairs evaluated. This results in approximately 75% of the cell’s reconstructed fragments being represented by at least one fragment *k*-mer pair. Although more distant paired *k*-mers tend to be more unique, and thus are better at the resolution of ambiguous branches in the DBG, increasing the distance between paired *k*-mers would lower the proportion of fragments represented by *k*-mer pairs. Therefore, it is important to balance the proportion of fragments represented by *k*-mer pairs and the distance between paired *k*-mers.

To avoid redundant storage, each fragment is screened against an assembled *k*-mers Bloom filter, which contains *k*-mers of previously reconstructed fragments, and the fragment *k*-mer pairs Bloom filter. If the fragment contains at least one new *k*-mer or *k*-mer pair, new *k*-mers and *k*-mer pairs are inserted into the corresponding Bloom filters. The fragment is then assigned to one of the strata according to its minimum *k*-mer count and its length (**Supplementary Fig. 6**). Fragments not consistent with reads are discarded, and their original paired-end reads are assigned to the strata for unconnected reads. This stratification of reconstructed transcript fragments provides a crude separation of fragment sequences of different expression levels. After all paired-end reads have been evaluated for a cell, both fragment *k*-mer pairs Bloom filter and the assembled *k*-mers Bloom filter are emptied in preparation for transcript fragment reconstruction of the next cell.

### Transcript sequence reconstruction

After fragment sequence reconstruction have been completed for all cells, transcript sequences can be reconstructed by extending each fragment sequence outward in both directions. To reconstruct transcript sequences for each cell, the DBG Bloom filter is emptied and repopulated with only *k*-mers along the cell’s reconstructed fragments. Emptying the DBG Bloom filter ensures that fragments are extended with *k*-mers specific to the corresponding cell.

Fragment sequences are retrieved from strata with decreasing *k*-mer counts because low-expression strata tend to be more enriched in sequencing errors and artifacts. Strata for long fragments are retrieved first, followed by the strata for short fragments, and finally strata for unconnected reads (**Supplementary Fig. 6**). Each fragment sequence is extended outward in both directions. The extension routine for each direction is similar to its counterpart in fragment sequence reconstruction, except that the scoring scheme here includes fragment *k*-mer pairs in addition to read *k*-mer pairs (**Supplementary Fig. 5** and **Supplementary Note 1**).

The reconstructed transcript sequences are evaluated for consistency with input reads and reconstructed fragments by scanning along the reconstructed transcript sequence for overlapping read *k*-mer pairs and fragment *k*-mer pairs. Segments of the transcript sequence without overlapping *k*-mer pairs are trimmed from the transcript sequence. This procedure ensures low number of mis-assembled transcripts.

### Software evaluations

We used RSEM^26^ to simulate 4 million paired-end reads of length 100 bp for 168 cells in the SMARTer scRNA-seq samples of mouse embryonic stem cells (ArrayExpress accession: E-MTAB-2600)^19^. The procedure for simulations is described in **Supplementary Note 3**. Benchmarking was performed with RNA-Bloom, Trans-ABySS, Trinity, and rnaSPAdes. The software versions and commands for each assembler are described in **Supplementary Note 4**. All assemblers were run using 48 threads on a machine with 48 HT-cores at 2.2 Ghz and 384 GB of RAM. Since the three bulk RNA-seq assemblers evaluated do not pool reads from multiple cells in their algorithms, the total run-time was the sum of those of the assemblies of individual cells. The number of full-length isoforms and missassemblies in each assembly were determined with rnaQUAST^27^, using the mouse reference genome GRCm38 and Ensembl version 91 annotations for measuring assembly sensitivity and correctness.

We calculated the quartiles of the expression levels measured on the transcripts per million (TPM) scale for all simulated isoforms in all 168 cells. We used these TPM quartiles to define four expression strata: (1) TPM < Q1, (2) Q1 ≤ TPM < Q2, (3) Q2 ≤ TPM < Q3, and (4) Q3 ≤ TPM. Every simulated isoform in each cell was assigned to one of the four strata. For each cell, we counted the number of isoforms reconstructed to ≥95%, as reported by rnaQUAST, within each stratum.

We used the dataset of 260 mouse embryonic stem cells (ArrayExpress accession: E-MTAB-2600)^19^ to investigate whether RNA-Bloom would scale to a much larger volume of input reads. Due to the input size, we used a maximum allowable Bloom filter FPR of 5% in RNA-Bloom to ensure low memory consumption.

### Fusion transcript discovery

We assembled the matching libraries of 62 K562 cells (SRA accession: SRP132313)^28^ from HiSeq and BGISEQ separately with RNA-Bloom. We supplied the assembled transcripts and the reads for each cell to PAVFinder for fusion detection. Similarly, we supplied the reads for each cell to STAR-Fusion for fusion detection. The software versions and commands for each method are described in **Supplementary Note 4**.

## Notes

https://github.com/bcgsc/RNA-Bloom

## REFERENCES

1. Song, Y., et al., Single-Cell Alternative Splicing Analysis with Expedition Reveals Splicing Dynamics during Neuron Differentiation. Mol Cell, 2017. 67(1): p. 148–161 e5.

2. Arzalluz-Luque, Á. and A. Conesa, Single-cell RNAseq for the study of isoforms-how is that possible? Genome Biol., 2018. 19(1): p. 110.

3. Conesa, A., et al., A survey of best practices for RNA-seq data analysis. Genome Biol., 2016. 17: p. 13.

4. Cole, C., et al., Tn5Prime, a Tn5 based 5’ capture method for single cell RNA-seq. Nucleic Acids Res., 2018. 46(10): p. e62.

5. Macosko, E.Z., et al., Highly Parallel Genome-wide Expression Profiling of Individual Cells Using Nanoliter Droplets. Cell, 2015. 161(5): p. 1202–1214.

6. Zheng, G.X.Y., et al., Massively parallel digital transcriptional profiling of single cells. Nat. Commun., 2017. 8: p. 14049.

7. Picelli, S., et al., Smart-seq2 for sensitive full-length transcriptome profiling in single cells. Nat. Methods, 2013. 10(11): p. 1096–1098.

8. Sheng, K., et al., Effective detection of variation in single-cell transcriptomes using MATQ-seq. Nat. Methods, 2017. 14(3): p. 267–270.

9. Canzar, S., et al., BASIC: BCR assembly from single cells. Bioinformatics, 2017. 33(3): p. 425–427.

10. Lindeman, I., et al., BraCeR: B-cell-receptor reconstruction and clonality inference from single-cell RNA-seq. Nat. Methods, 2018. 15(8): p. 563–565.

11. Rizzetto, S., et al., B-cell receptor reconstruction from single-cell RNA-seq with VDJPuzzle. Bioinformatics, 2018. 34(16): p. 2846–2847.

12. Robertson, G., et al., De novo assembly and analysis of RNA-seq data. Nat. Methods, 2010. 7(11): p. 909–912.

13. Chikhi, R. and G. Rizk, Space-efficient and exact de Bruijn graph representation based on a Bloom filter. Algorithms Mol Biol, 2013. 8(1): p. 22.

14. Jackman, S.D., et al., ABySS 2.0: resource-efficient assembly of large genomes using a Bloom filter. Genome Res, 2017. 27(5): p. 768–777.

15. Bloom, B.H., Space/time trade-offs in hash coding with allowable errors. Commun. ACM, 1970. 13(7): p. 422–426.

16. Bray, N.L., et al., Near-optimal probabilistic RNA-seq quantification. Nat Biotechnol, 2016. 34(5): p. 525–7.

17. Grabherr, M.G., et al., Full-length transcriptome assembly from RNA-Seq data without a reference genome. Nat Biotechnol, 2011. 29(7): p. 644–52.

18. Bushmanova, E., et al., rnaSPAdes: a de novo transcriptome assembler and its application to RNA-Seq data. bioRxiv, 2018.

19. Kolodziejczyk, A.A., et al., Single Cell RNA-Sequencing of Pluripotent States Unlocks Modular Transcriptional Variation. Cell Stem Cell, 2015. 17(4): p. 471–85.

20. Chiu, R., et al., TAP: a targeted clinical genomics pipeline for detecting transcript variants using RNA-seq data. BMC Med. Genomics, 2018. 11(1): p. 79.

21. Haas, B., et al., STAR-Fusion: Fast and Accurate Fusion Transcript Detection from RNA-Seq. Bioinformatics, 2017.

22. Mohamadi, H., et al., ntHash: recursive nucleotide hashing. Bioinformatics, 2016. 32(22): p. 3492–3494.

23. Mohamadi, H., H. Khan, and I. Birol, ntCard: a streaming algorithm for cardinality estimation in genomics data. Bioinformatics, 2017. 33(9): p. 1324–1330.

24. Birol, I., et al., Spaced Seed Data Structures for De Novo Assembly. Int J Genomics, 2015. 2015: p. 196591.

25. Song, L. and L. Florea, Rcorrector: efficient and accurate error correction for Illumina RNA-seq reads. Gigascience, 2015. 4: p. 48.

26. Li, B. and C.N. Dewey, RSEM: accurate transcript quantification from RNA-Seq data with or without a reference genome. BMC Bioinformatics, 2011. 12: p. 323.

27. Bushmanova, E., et al., rnaQUAST: a quality assessment tool for de novo transcriptome assemblies. Bioinformatics, 2016. 32(14): p. 2210–2.

28. Natarajan, K.N., et al., Comparative analysis of sequencing technologies for single-cell transcriptomics. Genome Biol, 2019. 20(1): p. 70.

