## Supplementary information for "RNA-Bloom provides lightweight reference-free transcriptome assembly for single cells"

**Supplementary Figure 1.** Pooling of reads from multiple cells. 2

**Supplementary Figure 2.** Number of full-length isoforms reconstructed by each method. 3

**Supplementary Figure 3.** Paired *k*-mers along reads and reconstructed fragments. 4

**Supplementary Figure 4.** Error correction in paired-end reads. 5

**Supplementary Figure 5.** *K*-mer pairs spanning candidate extensions. 6

**Supplementary Figure 6.** Stratification of reconstructed fragments and unconnected reads. 7

**Supplementary Note 1.** Heuristic score for selecting the best candidate extension. 8

**Supplementary Note 2.** Cell-specificity of pooled assembly. 8

**Supplementary Note 3.** Commands for running various programs. 9

**Supplementary Table 1.** Known gene fusions in the K562 cell line. 11

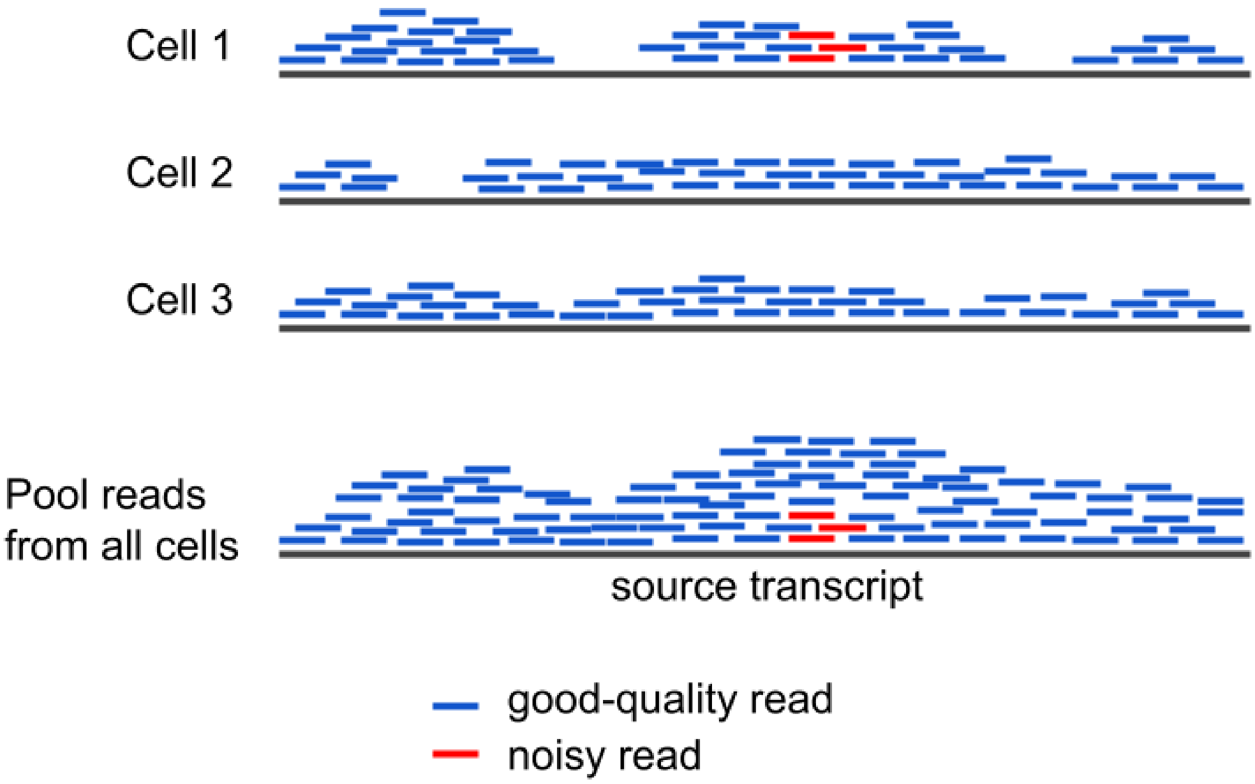


**Supplementary Figure 1.** Pooling of reads from multiple cells.

Pooling reads from multiple cells improves transcript sequence reconstruction by filling in sequencing gaps and increasing the signal-to-noise ratio.

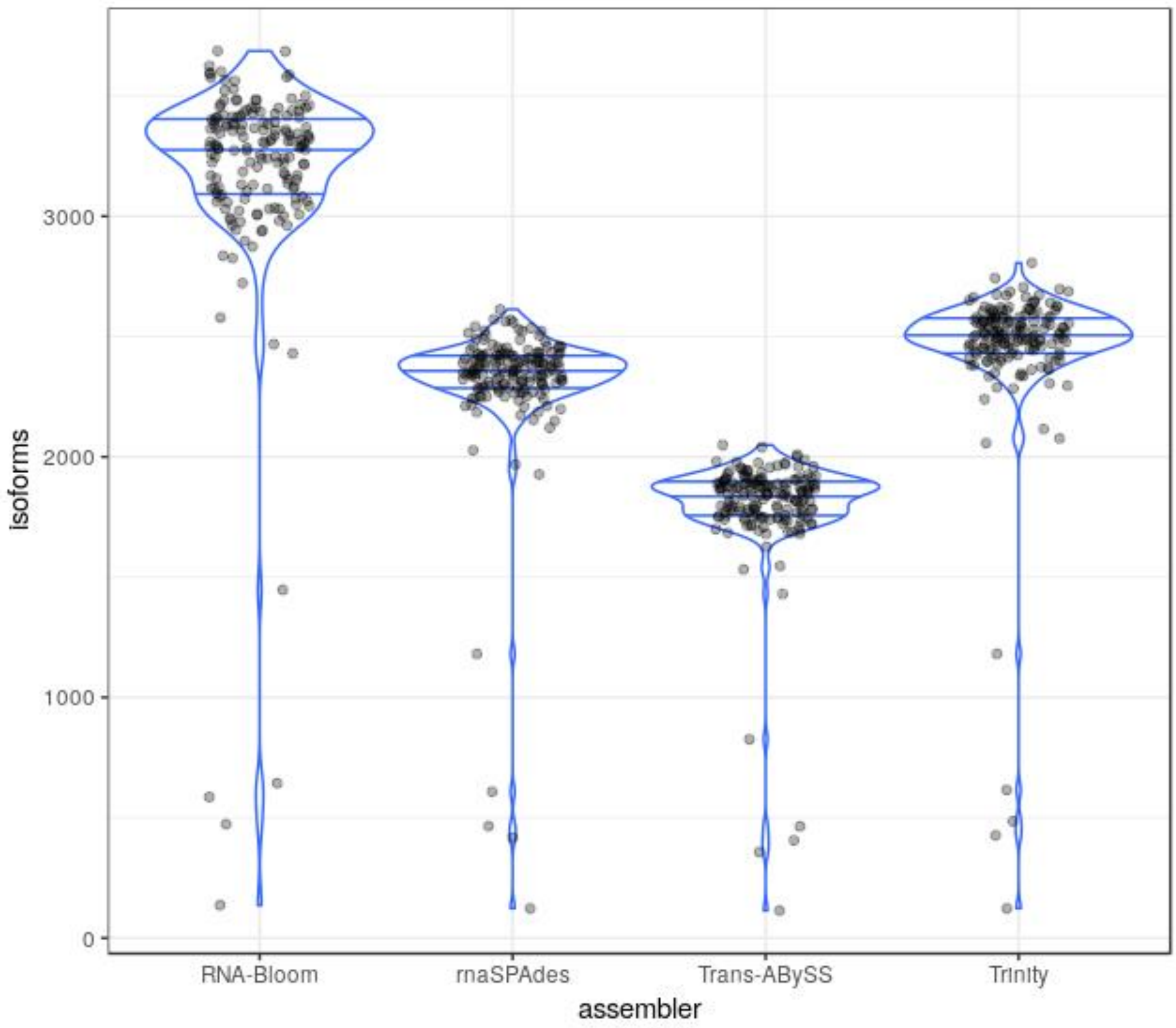


**Supplementary Figure 2.** Number of full-length isoforms reconstructed by each method.

The isoform reconstruction of each method is measured by rnaQUAST. An isoform is reconstructed to "full length" if at least 95% of its annotated length is spanned by one assembled contig.

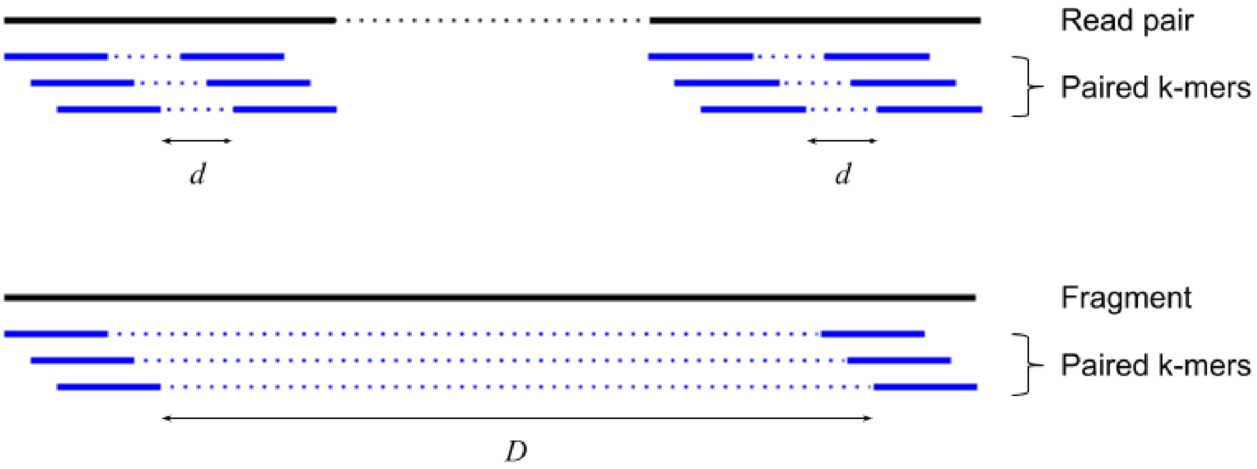


**Supplementary Figure 3.** Paired *k*-mers along reads and reconstructed fragments.

Paired *k*-mers are sparse representations of the sequences from where they were derived. This is analogous to a read pair being a sparse representation of its underlying fragment. The distance between read paired *k*-mers (*d*) is set based on the read length, and it is the same for all cells. The distance between fragment paired *k*-mers (*D*) is set based on the first quartile fragment length for each cell.

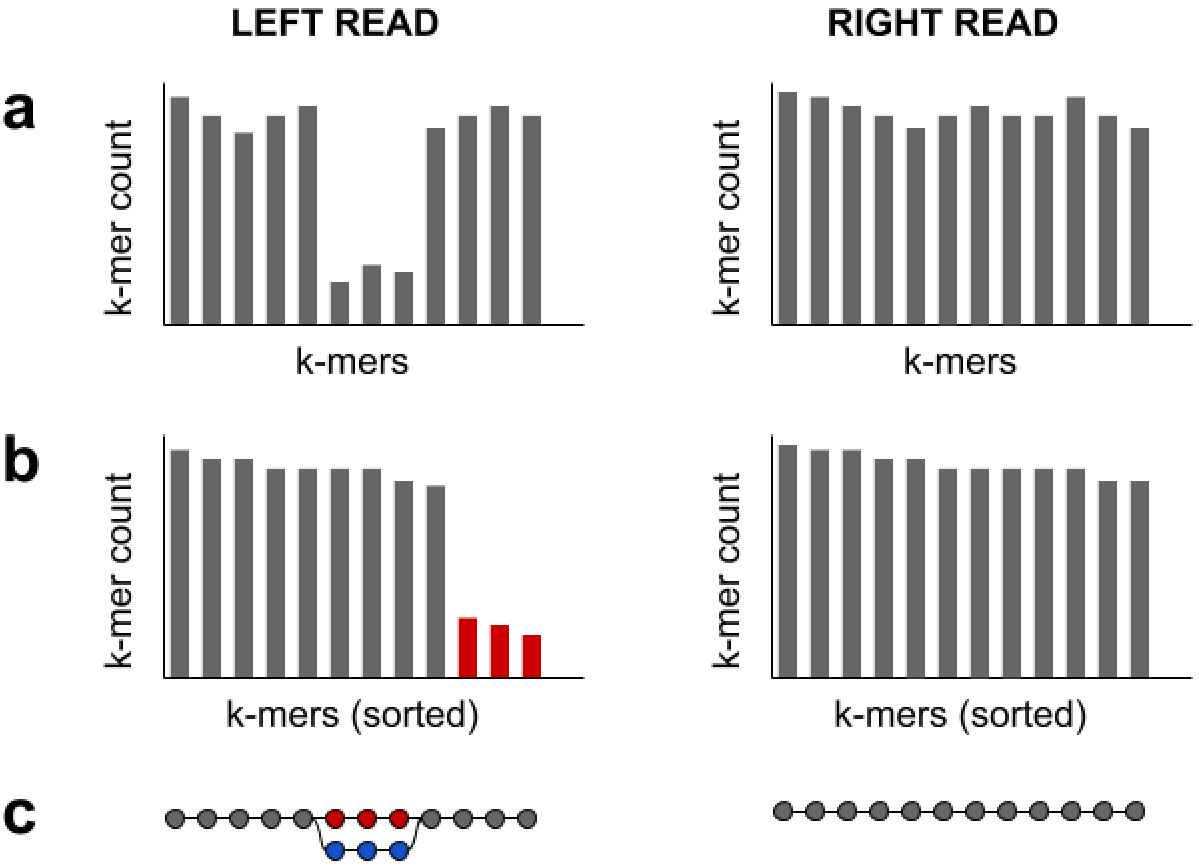


**Supplementary Figure 4.** Error correction in paired-end reads.

(**a**) Errors in paired-end reads are corrected based on *k*-mer counts.
(**b**) A local *k*-mer count threshold is determined for each read individually. This is done by sorting the *k*-mer counts in descending order, and then looking for the first sharp decrease (defined as ≥ 50%), where the *k*-mer count before the sharp decrease is set as the local *k*-mer count threshold. If there is no sharp decrease in *k*-mer counts, then the read would not be corrected. If both reads have their own local thresholds, then the lower threshold is set for both reads to account for low-expressed alternative isoforms.
(**c**) *K*-mers with counts lower than the threshold are marked (ie. red nodes) for replacement with an alternative branch (ie. blue nodes) in the DBG that has a higher median *k*-mer count, similar length, and high sequence identity (default ≥ 90%).

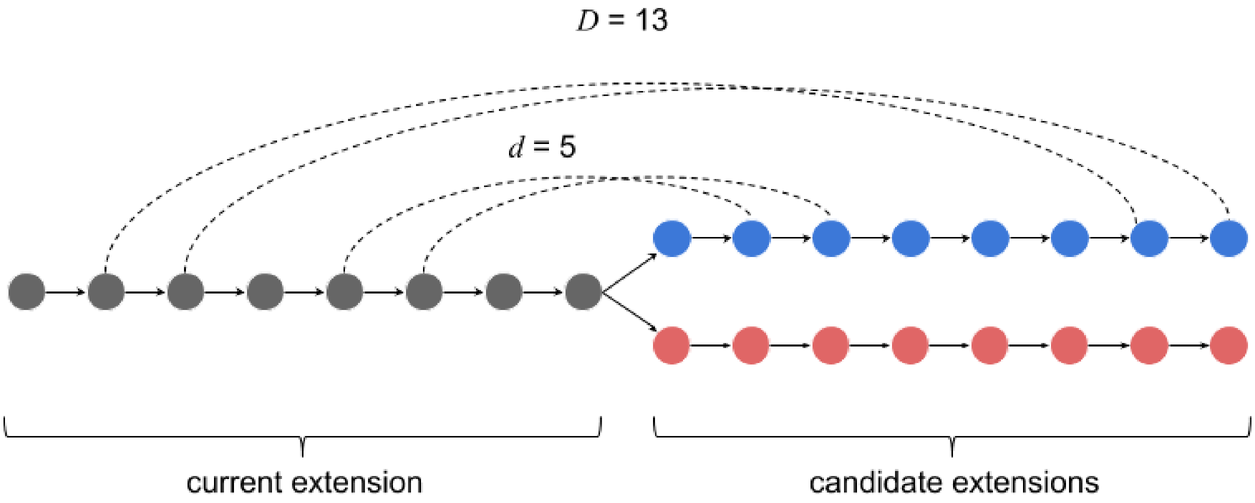


**Supplementary Figure 5.** *K*-mer pairs spanning candidate extensions.

When an extension reaches a fork in the DBG, *k*-mer pairs from reads and fragments are used to identify the best candidate extension from the branching point. In this example, read paired *k*-mers have a distance, $d = 5$, and fragment paired *k*-mers have a distance, $D = 13$. The blue branch is a better candidate extension compared to the red branch because it has two supporting read *k*-mer pairs and two supporting fragment *k*-mer pairs.

| Minimum k-mer count ($c$) | Fragment length ($L$) | | Unconnected reads |
|---|---|---|---|
| | $L \geq$ Q1 “Long fragments” | $L <$ Q1 “Short fragments” | |
| $100{,}000 \leq c$ | I | VII | XIII |
| $10{,}000 \leq c < 100{,}000$ | II | VIII | XIV |
| $1{,}000 \leq c < 10{,}000$ | III | IX | XV |
| $100 \leq c < 1{,}000$ | IV | X | XVI |
| $10 \leq c < 100$ | V | XI | XVII |
| $1 < c < 10$ | VI | XII | XVIII |
| $c = 1$ | XIX | XX | XXI |

**Supplementary Figure 6.** Stratification of reconstructed fragments and unconnected reads.

Reconstructed fragments are stratified according to their length and minimum *k*-mer count. Since the fragment paired *k*-mers distance is set based on the first quartile fragment length, “long” fragments have at least one fragment *k*-mer pair while “short” fragments do not. Unconnected reads are stratified according to their minimum *k*-mer count only. The stratum IDs (roman numerals in the table) denote the order in which the fragments are retrieved for extension into transcript sequences. Sequences with singleton *k*-mers (ie. $c$ = 1) are extended lastly because they often have sequencing artifacts.

### **Supplementary Note 1.** Heuristic score for selecting the best candidate extension.

A heuristic score ($H$) can be calculated to select the best candidate extension based on the following 4 parameters:

1. median $k$-mer count of extension ($c$)
2. number of supporting read $k$-mer pairs ($r$)
3. number of supporting fragment $k$-mer pairs ($f$)
4. number of $k$-mers in extension ($l$)

During fragment sequence reconstruction:

$$H = c\, r / l$$

During transcript sequence reconstruction:

$$H = c\, (r + f) / l$$

$$H = 0 \text{ if } r = 0 \text{ or } f = 0.$$

### **Supplementary Note 2.** Cell-specificity of pooled assembly.

The measure of cell-specificity ($S$) of a pooled assembly is derived from the mathematical equation for specificity, ie.

$$S = TN / (TN + FP)$$

where:
TN is the number of isoforms unique to other cells and are not reconstructed for this cell
FP is the number of isoforms unique to other cells but reconstructed for this cell

**Supplementary Note 3.** Commands for running various programs.

**RSEM v1.3.1:**

*(prepare reference)*

```
rsem-prepare-reference --gtf GRCm38.ensembl91.gtf --star \
      --star-path STAR-2.6.1a/bin/Linux_x86_64 \
      GRCm38.dna.primary_assembly.fa mouse_ref
```

*(quantify transcript expression for each cell)*

```
rsem-calculate-expression -p 24 --paired-end --star \
      --star-path STAR-2.6.1a/bin/Linux_x86_64 \
      --estimate-rspd --append-names --calc-ci --single-cell-prior \
      --output-genome-bam LEFT.fastq RIGHT.fastq mouse_ref NAME
```

*(simulate scRNA-seq reads for each cell)*

```
rsem-simulate-reads mouse_ref NAME.stat/NAME.model \
      NAME.isoforms.results 0.2 2000000 OUTDIR
```

**RNA-Bloom v1.0.0:**

*(assemble transcripts for all cells)*

```
java -Xmx48g -jar RNA-Bloom.jar -ntcard -fpr 0.005 -k 25 -extend \
      -stratum 01 -t 48 -pool READSLIST.txt -revcomp-right -outdir OUTDIR
```

**Trans-ABySS v2.0.0:**

*(assemble transcripts for each cell separately)*

```
transabyss --pe LEFT.fastq RIGHT.fastq --threads 48 -k 25 -c 1 --length 200 \
      --outdir OUTDIR
```

**Trinity v2.5.1:**

*(assemble transcripts for each cell separately)*

```
Trinity --seqType fq --left LEFT.fastq --right RIGHT.fastq --CPU 48 \
      --max_memory 4G --output OUTDIR
```

**rnaSPAdes v3.11.1:**

*(assemble transcripts for each cell separately)*

```
rnaspades.py -t 48 -1 LEFT.fastq -2 RIGHT.fastq -o OUTDIR
```

**rnaQUAST v1.5.2:**

(*evaluate assemblies for each cell separately*)

```
rnaQUAST.py -r GRCm38.dna.primary_assembly.fa --gene_db GRCm38.ensembl91.db \
      --disable_infer_genes --disable_infer_transcripts -t 12 \
      --blat --no_plots \
      -c RNABLOOM.fasta TRANSABYSS.fasta TRINITY.fasta RNASPADES.fasta \
      -l RNA-Bloom Trans-ABySS Trinity rnaSPAdes \
      -o OUTDIR
```

**GMAP version 2019-05-12:**

(*align assembly to reference genome*)

```
gmap -d hg38 -D hg38_index_dir ASSEMBLY.fasta -t 24 -f samse -n 0 | \
      samtools view -bhS - -o C2G.bam
```

**BWA v0.7.12 (r1039):**

*(align reads to assembly for each cell)*

```
bwa index ASSEMBLY.fasta
bwa mem -t 24 ASSEMBLY.fasta LEFT.fastq RIGHT.fastq | \
      samtools view -bhS - | \
      samtools sort -m 20G - -o R2C.bam
samtools index R2C.bam
```

*(align assembly to reference transcriptome for each cell)*

```
bwa mem -t 24 hg38_gencode_v26_transcripts.fa ASSEMBLY.fasta | \
      samtools view -bhS -o C2T.bam
```

**PAVFinder v1.5.0:**

(*detect fusions from assembly and alignments for each cell*)

```
pavfinder fusion \
      --transcripts_fasta hg38_gencode_v26_transcripts.fa \
      --genome_index hg38_index_dir hg38 \
      --only_fusions --include_non_exon_bound_fusion --min_support 1 \
      --gbam C2G.bam --tbam C2T.bam --r2c R2C.bam \
      ASSEMBLY.fasta hg38_gencode_v26.gtf hg38_genome.fa OUTDIR
```

**STAR-Fusion v1.1.0 (using STAR v2.5.3a):**

(*detect fusions from alignments for each cell*)

```
STAR-Fusion --genome_lib_dir star_reference_index_dir --CPU 24 \
      --left_fq LEFT.fastq --right_fq RIGHT.fastq --output_dir OUTDIR
```

| **Gene fusion** | **References** |
|---|---|
| *ABL1:BCR* | ● https://depmap.org/portal/cell_line/K562_HAEMATOPOIETIC_AND_LYMPHOID_TISSUE?tab=fusion<br>● Genome Biol. 2009;10(10):R115. doi: 10.1186/gb-2009-10-10-r115. |
| *BAG6:SLC44A4* | ● https://depmap.org/portal/cell_line/K562_HAEMATOPOIETIC_AND_LYMPHOID_TISSUE?tab=fusion |
| *NUP214:XKR3* | ● Genome Biol. 2009;10(10):R115. doi: 10.1186/gb-2009-10-10-r115. |
| *CBX5:GTSF1* | ● https://ccsm.uth.edu/FusionGDB/gene_search_result.cgi?page=page&type=quick_search&quick_search=5717 |
| *C16orf87:ORC6* | ● https://depmap.org/portal/cell_line/K562_HAEMATOPOIETIC_AND_LYMPHOID_TISSUE?tab=fusion |

**Supplementary Table 1.** Known gene fusions in the K562 cell line.
